# Castes and developmental stages of the harvester ant *P. californicus* differ in genome-wide and gene specific DNA methylation

**DOI:** 10.1101/2025.03.13.642958

**Authors:** Tania Chavarria-Pizarro, Mohammed Errbii, Janina Rinke, Lukas Schrader, Jürgen Gadau

## Abstract

DNA methylation in social insects has been proposed as a key mechanism underlying both reproductive and non-reproductive caste determination by regulating gene expression in response to environmental cues. So far, studies on DNA methylation in social insects have yielded mixed results, with some studies finding no effect on caste determination or worker behaviour while others have reported such effects. In order to link DNA methylation to either process one first needs to show variation in DNA methylation between castes and developmental stages. This study determined cytosin methylation in a CpG context (C^m^pG) in the harvester ant *Pogonomyrmex californicus* using ONT (Oxford Nanopore Sequencing) sequencing in larvae, pupae, workers and queens, moreover we compared accuracy of methylation with the gold standard WGBS (Whole Genome Bisulfite Sequencing). Methylation sites were highly correlated between WGBS and ONT (r^2^=0.8). Overall, the *P. californicus* genome showed low levels of methylation (3%) which is comparable to what has been found in other Hymenopterans. DNA methylation was significantly increased in gene bodies and in particular exons and showed a significant decrease at the transcription start site and promotor region. Introns, intergenic regions and transposable elements have the lowest levels of DNA methylation. Gene body methylation was positively correlated with gene expression in queens, which is in accordance with other studies and the opposite of what is found in vertebrate studies. Both castes and developmental stages showed significant variation and differences in gene body methylation frequencies. Furthermore, castes and stages which have a high metabolic rate (queens and pupae) show higher DNA methylation, suggesting plasticity and an active role of DNA methylation in gene regulation and in extension caste regulation.

## Introduction

Epigenetics is the study of phenotypic changes in organisms generated by alterations of gene expression rather than changes to the DNA sequence (Holliday 1975, Jablonka & Lamb 2002, Richards 2006, Bird 2007, Dean & Maggert 2015). Epigenetic regulation includes DNA methylation, histone modification, chromatin remodelling, and ncRNA (Grant-Downton & Dickinson 2005, Berger 2007). The ensemble of all epigenetic changes affecting the genome, also known as the epigenome, has been shown to have heritable components (Murrel et al. 2005, Kungulovski & Jeltsch 2016). Among epigenetic mechanisms, DNA methylation is arguably the most extensively studied epigenetic mechanism (Duncan et al. 2014). In particular, the addition of a methyl group to the fifth cytosine’s carbon (5mC) is the most common and well-studied form of DNA methylation and is found across a wide range of organisms including bacteria, plants, fungi, invertebrates, and vertebrates (Feng et al. 2010, Suzuki & Bird 2008, Duncan et al. 2014, Kungulovski & Jeltsch 2016). Where it has also been shown to regulate gene expression. However, its functions, biological characteristics, and genomic distribution are unique for each taxonomic lineage and varies considerable within each lineage (Colot & Rossignol 1999). For instance, studies in vertebrates demonstrated that methylation of cytosines within a CpG context (5mCpG), reduced transcription of the methylated genes by blocking the binding of transcription factors to the DNA (Lyko 2018). DNA methylation as control of the TEs activity known as epigenetic silencing has been proposed as an important process to repress the potentially detrimental effects of TE expression (Bestor 1990). DNA methylation especially represses TEs in many higher eukaryotes (e.g. Deniz et al. 2019, Qiu et al. 2023).

In contrast, the role of DNA methylation in insects is less clear, as it varies widely across insects with some species showing no methylation (*Drosophila melanogaster*) while others (*Blattella germanica*) exhibit considerable levels of DNA and TEs methylation (Lewis et al. 2020). DNA methylation in insects is predominantly found in coding regions associated with gene bodies of actively expressed genes (Suzuki & Bird 2008, Lyko et al. 2010, Zemach et al. 2010, Bonasio et al. 2012). Furthermore, DNA methylation tends to occur within exons of highly conserved housekeeping genes essential for cellular functions (Sarda et al. 2012, Dimond & Roberts 2016) and has been linked to an upregulation of genes involved in several important biological functions, such as reproduction, embryogenesis, and development (Kucharski et al. 2008, Lyko et al. 2010, Shi et al. 2013, Yang et al. 2017, Kay, Skowronski & Hunt 2017).

In social insects, DNA methylation has been proposed as a key mechanism underlying caste determination by regulating gene expression. This is because they are hallmarks of phenotypic plasticity where reproductive, morphological and behavioural castes can develop from the same genome (Kucharski et al. 2008, Herb et al. 2012, Yan, Bonasio, Simola, & Berger 2015). Hence, social insects are ideal models to investigate the function of DNA methylation in caste determination and more generally, phenotypic plasticity (Lyko & Maleszka 2011). So far, studies on the role of DNA methylation in social insects have yielded mixed results, with some studies finding no effect on caste determination or worker behavior (Patalano et al. 2015, Libbrecht et al. 2016, Oldroyd & Yagound 2021), while others have reported such effects (Lyko & Maleszka 2011, Morandin et al. 2019). Morandin et al. (2019) found differences in expression and methylation profiles between workers and queens at different life stages, as well as some overlap between DNA methylation and expression in brains of the ant *Formica exsecta*. In contrast, Cardoso-Júnior et al. (2021) found no significant differences between the methylomes of workers from queen-right and queen-less colonies of honeybees, moreover differentially methylated regions were not associated with differential gene expression and methylation patterns were highly similar between brain and ovary tissues.

As one explanation for the inconsistent results of these different studies Morandin et al. (2019) suggested that factors such as a lack of biological replicates and appropriate methods to detect DNA methylation could explain those inconsistencies. Additionally, Wedd et al. (2016) showed that DNA methylation can be allele specific and its link to expression is only measurable at specific time points during development, when expression is in high demand (Wedd et al. 2016). Overall, the function and role of DNA methylation in insects is still not well understood and it is unclear whether results from one species can be generalized.

Several techniques have been developed to detect and measure epigenetic marks in natural populations, particularly DNA methylation (Husby et al. 2022). One widely used method is Whole Genome Bisulfite Sequencing (WGBS), where the DNA is treated with sodium bisulfite, converting unmethylated cytosines into uracils while leaving methylated cytosines unchanged. Oxford Nanopore Technologies (ONT) sequencing is another method gaining popularity because it does not need chemical treatment with sodium bisulfite. Although, both methods are highly effective, they also come with limitations. For example, bisulfite conversion degrades DNA due to depyrimidination of unmethylated cytosines and typically yields low sequencing coverage across GC-rich regions (Feng et al. 2020, Olova et al. 2018, Guanzon et al. 2024). The bias on methylation calls with ONT method could result from sequence context, or due to imbalances in coverage, like in shorter reads sequencing methods (Guanzon et al. 2024). In addition, a systematic comparison of ONT methylation callers revealed that readout biases are largely attributable to the choice of methylation caller (Yueng et al. 2021, Guanzon et al. 2024), nonetheless this bias disappears when the coverage is higher than 10x (Guanzon et al. 2024). High correlations have previously been found between ONT and WGBS, especially when methylation frequencies were averaged over large genomic windows (1-100 kb) corresponding to gene sizes (Perez et al. 2023). Faulk (2023) convincingly showed that even low coverage ONT sequencing (genome skimming) predicted reliably global DNA methylation equally well as WGBS and had an average mapping rate of 97 %. This study also demonstrated that ONT was very successful in predicting transposon methylation due to its long reads.

Here, we compared genome wide DNA methylation in the harvester ant *Pogonomyrmex californicus* using both WGBS and ONT techniques to conduct the first epigenetic survey in this species. Our objectives were 1) Evaluate the accuracy and correlation between ONT sequencing and WGBS, using identical DNA pools (pupae) 2) Assess methylation patterns along functional different genome regions, TEs and TEs islands, to shed light on the function and role of DNA methylation in insects 3) Compare specific DNA methylation patterns between three different life stages (larva, pupae, adult) and two female castes (queens and workers), to elucidate a putative link between DNA methylation and caste evaluating its potential to explain social variation and behavioural plasticity in this species 4) Evaluate if DNA methylation correlates with gene expression in queens.

## Methods

### Specimen collection and samples

We used different life stages of *Pogonomyrmex californicus* (larva, pupae, and workers) from queenright colonies kept at the Institute for Evolution and Biodiversity in Münster. Note, since these colonies do not produce sexuals, males or queens, we can be sure that both larvae and pupae would have developed into workers. Those laboratory colonies were established from founding queens of *P. californicus* collected in 2018 and 2019 at Pine Valley (32.822 N, − 116.528 W; pleometrotic population, P-population) and Lake Henshaw resort (33.232 N, − 116.760 W; haplometrotic population, H-population), California, USA.

We also used founding queens collected directly after their nuptial flight (Salt River, Phoenix Arizona 2023 (33.55034 N, − 111.64453 W). We prepared ONT libraries for ten queens, two pools of 4 workers, two pools of 4 pupae, and two pools of 4 larvae each (Table 1). Each pool was generated from individuals of the same colony. In addition, we used the same pupal pools to generate two WGBS libraries to directly compare the results of ONT sequencing and WGBS.

**Table 1.**
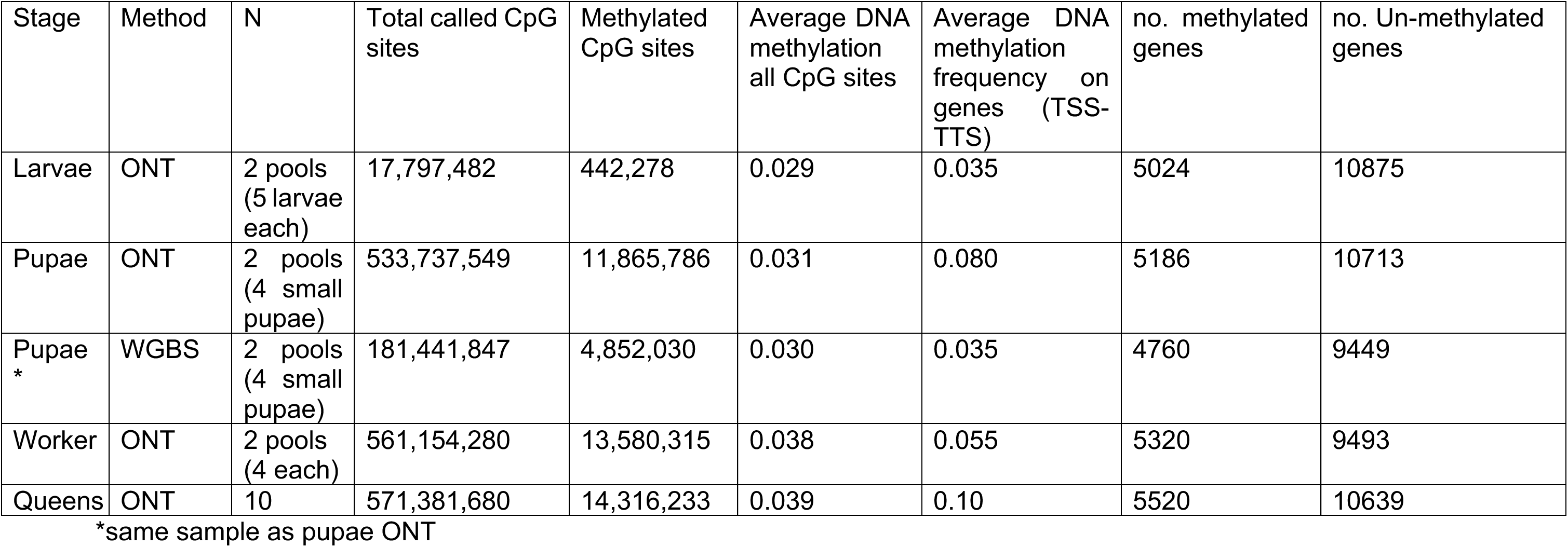
Nanopore ONT and WGBS sequencing – methylation data per stage and caste. Average methylation was calculated as the average CG gene methylation across all genes. We calculated average DNA methylation for each stage and caste separately, as methylation levels differ between these categories. A gene was considered methylated if had a significantly higher proportion of methylated cytosines than the genome-wide methylation average in each stage or caste. This was performed using a Binomial test and p-values were corrected for multiple tests.

### DNA isolation and sequencing

DNA was extracted from whole bodies of frozen or ethanol-preserved individuals of *P. californicus* with the Qiagen DNeasy Blood and Tissue kit. A Qubit BR assay was used to assess DNA quantity, followed by 1 % agarose gel electrophoresis to confirm the presence of high molecular weight DNA (>10 kb). We prepared DNA libraries for sequencing using the Oxford Nanopore Technologies (ONT) MinION platform according to the user manual. Following the manufacturer guidelines we used PCR-free Ligation Sequencing Kit (PCR-free ONT Ligation Sequencing Kit with the Native Barcoding Expansion Kit (SQK-LSK109 and EXPNBD103), and transposase-based ONT Rapid Sequencing Kit (SQK-RAD004). Sequencing was done on FLO-MIN 106D R9.4 flow cells, following the manufacturer’s 1D native barcoding gDNA protocol. Each library was run on a separate FLO-MIN 106D R9.4 flow cell. Basecalling fast 5 and demultiplexing was performed with ont-guppy/6.4.8-CUDA-11.7.0 using the appropriate base calling model (dna_r9.4.1_450bps_hac.cfg) and default parameters (More details in https://github.com/TaniaChP79/P.cal-methylome).

### Genome Assemblies and Annotations

We used the newest genome assembly and annotation for *P. californicus* (Pcal.3.1) (Errbii et al. unpublished) based on combined ONT long-read sequencing and 10 Å∼ sequencing. The annotation included 15,899 protein-coding genes and had a TE proportion of 22.79 % (Errbii et al. unpublished).

### CpG Methylation Calling

The Nanopolish v0.13.3 (Simpson et al. 2017) pipeline was used with default parameters to detect CpG methylation in ONT data. Nanopolish is computationally efficient and has previously been used in methylation studies using ONT sequencing data (Liu et al. 2021, Yuen et al. 2021). Briefly, the Nanopolish software uses a pre-trained hidden Markov model to assign methylation log-likelihood ratios (LLRs) to all CpGs within a 10 bp window. First, Nanopolish indexes the nanopore reads and then maps these onto the reference genome of *P. californicus* (Pcal 3.1) using minimap2 v. 2.14 (Li 2018). After alignment the Nanopolish software analyses short CpG motif (11– 34 bp) k-mer sequences and separates 5-methylcytosine from unmethylated cytosines based on signal disruptions in the raw ONT FAST5 sequence data. After that, the program calculates log-likelihood ratios for base modifications of each read, where positive values indicate support for modification (methylation). We used the helper script *calculate_methylation_frequency.py* (Simpson et al. 2017) to convert the Nanopolish output into methylation frequency by genomic coordinates. The methylation frequency in Nanopolish is calculated as the proportion of reads that show evidence of methylation at a particular site relative to all reads covering that site. This provides a measure of how often a cytosine is methylated in the sampled population of molecules (Simpson et al. 2017).

The methylation calls were then subject to filtering using a minimum read count of 10 reads per CpG motif. We excluded genome sequences with methylation calls of less than 10% of CpGs in the sequence as suggested by Perez et al (2023). Additionally, a bed file with all CpGs for the genome was generated and mapped to the respective methylation annotations (call coverage and frequency) with the *bedtools map* function (Quinlan & Hall 2010), both for the transcriptome annotation and the TE annotation (Errbii et al. unpublished).

### Whole Genome Bisulfite sequencing

The WGBS sequencing libraries were prepared by Novogene (Munich, Germany). In short, paired-end 150 bp bisulfite libraries were sequenced on the Illumina NovaSeq 6000 platform to a total of 35 million paired-end reads (nuclear coverage >20X). Reads were trimmed with trimmomatic v0.30 (Bolger, Lohse & Usadel 2014) (parameters: leading = 10, trailing = 10, minlen = 50) and processed with bismark v0.22.3 (Krueger & Andrews 2011) to compute per-base-pair methylation frequencies. For WGBS data, we calculated methylation sites of pupae using the R package *methylKit v1.15.3* and R version 4.5.1 (Akalin et al. 2012). The percentage of methylated cytosines was calculated at a given site from the methylation ratios created by the software BSSeeker2 and complementary CpG dinucleotides were merged.

### Comparison between ONT and WGBS

We compared DNA methylation calls derived from ONT sequencing (sequencing depth > 40x coverage) with whole-genome bisulfite sequencing (WGBS, sequencing depth > 20x coverage), from the same extracted DNA from pooled pupae (n=2) which were used for the ONT sequencing. CpG motifs uniquely called either with Bismark or Nanopolish were further extracted using *bedtools subtract* (Quinlan & Hall 2010). The per-CpG motif methylation rates detected by ONT and WGBS were correlated using a Pearson correlation.

### TE annotation and inference of TE islands

TEs were annotated with RepeatMasker 4.1.7, using the newest assembly (Pcal 3.1). We created bed files of the TE and exon content across these pre-defined windows to calculate TE islands. TE distribution was plotted for 50 kb, 200 kb, and 500 kb genomic windows using *bedtools makewindows*, *sortBed* and *bedmap* from the bedtools package (Fig S1.). TE islands were defined for the 16 chromosomes across 200 kb windows using a custom script in R, applying an automated model to call TE islands with the packages *EnvCpt, Rbeast, gdata, zoo,* and *cowplot*. Parameters were set with a cutoff of 0.2 for the environment mean drop to call TE islands and a penalty of 40 for the environment mean model. All visualizations and analyses were done in R. We compared average methylation frequency on the TE islands with the rest of the genome using a Wilcoxon rank sum test.

### Inferences of gene body methylation

To exclude low-coverage data we set a commonly used threshold of at least 10 methylation sites per k-mer per sample (Perez et al. 2023).

For our analysis of average DNA methylation across all annotated genes, we located the methylation context within a window of 4 kb upstream of the transcription start site (TSS) and a window of 4 kb downstream of the transcription termination site (TTS). Then, we looped over all genes to compute average methylation proportions on genomic windows (4kb) upstream, downstream, and inside of genes. We computed the average CG gene methylation across all genes and performed a binomial test (Takuno & Gaut 2012), to assess whether coding regions had a significantly higher proportion of methylated cytosines than the genome-wide background level of non-coding regions, as methylation levels differ by developmental stage and caste, we performed binomial test separately by stage and castes (Bewick et al. 2016, Muyle et al. 2021). This was performed for each cytosine in a CpG context and p-values were corrected for multiple testing using Benjamin and Hochberg (1995) correction for each stage separately. Then we classified gene methylation into two categories: 1) Gene Body Methylated (GbM), if the adjusted p-value for CpG methylation was higher than expected by chance and 2) Un-methylated (UM), if the adjusted p-value for CG methylation was lower than expected by chance.

Finally, a GO term enrichment analysis was performed for all genes with significant gene-body methylation, using the *topGO* package (2.56) (Alexa & Rahnenfuhrer 2024), in R (4.5.0.) with a p-value < 0.05 representing significantly enriched GO terms.

### Inferences of gene promoter methylation

To study methylation of promoters, we used a window of 200 bp upstream of the TSS. We computed an average proportion of methylated promoter CpG sites and compared the resulting values to the genome average using a similar strategy to the gene body methylation described above (Bewick et al. 2016, Muyle et al. 2021). First, we used a binomial test (Takuno & Gaut 2012) and corrected obtained p-values for multiple tests using Benjamin and Hochberg (1995) (see section above for more details). Then, we classified gene promoter methylation into two categories 1) Promoter Gene Methylated (PGM) if adjusted p-values were higher than expected and 2) promote Un-methylated (PUM) if adjusted p-values were lower than expected. We did this analysis for each gene promoter region of each life stage. A promoter was considered methylated if the value was above 3 % for all the stages and castes. Similarly, a GO enrichment analysis was performed for all methylated promoters.

### Gene expression analysis

We used a *P. californicus* transcriptomes generated from 22 founding queens of *P. californicus* collected in 2018 and 2019 at Pine Valley (32.822 N, − 116.528 W; pleometrotic population, P-population) and Lake Henshaw resort (33.232 N, − 116.760 W; haplometrotic population, H-population), California, USA. Raw RNA-sequence reads were produced using the same process as Helmkampf et al. (2016). The reads were trimmed to exclude low-quality reads using Trimmomatic (v.0.39) (Bolger, Lohse & Usadel 2014) (parameters: leading = 10, trailing = 10, minlen = 50). The software HISAT2 (v.2.2.1) (Kim, Langmead & Salzberg 2015) was used to map the reads to the *P. californicus* reference genome (Erbii et al. unpublished). FeatureCounts (v.2.0.1) (Liao, Smyth & Shi 2014) was used to calculated raw counts per gene. Then we converted the raw counts to FPKM (Fragments Per Kilobase Million) for our gene expression data, calculating the length of each gene in base pairs and converting the gene length from base pairs to kilobases (kb) by dividing it by 1,000. Furthermore, we calculated the total mapped reads summing up raw counts of all genes to get the total number of reads mapped for each sample (total count across all genes). We then applied the FPKM formula as:

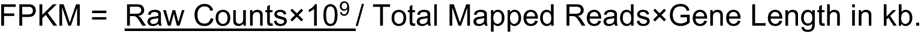

### Dnmt1 and Dnmt3 methylation frequency along life stages and castes

We calculated methylation frequency along life stages and castes of the maintenance (Dnmt1) and de novo (Dnmt3) DNA methyltransferases using the data generated with ONT.

## Results

The methylome of *P. californicus* was successfully characterized using a combination of long (ONT) and short reads (WGBS). Larvae, pupae, workers, and individual queens were analysed using ONT sequencing. Additionally, the same two pupae pools were also analysed by WGBS. This allowed a direct correlation of ONT sequencing and WGBS comparing overlap and efficiency of both techniques (Table 1). As a backbone for all analyses, we used a recently updated chromosome-level genome assembly and its annotation for *P. californicus* (Pcal3.1, Errbii et al. unpublished).

WGBS is still considered the gold standard to determine DNA methylation. We found that around 3% cytosines in a CpG context were methylated using both methods (ONT and WGBS). However, WGBS found consistently fewer methylated sites in all genomic contexts (Fig. 1 B, gene body and D, intergenic sites) which could be explained in part by the lower coverage of WGBS (ONT sequencing: > 40X coverage; 99 % CpGs sites called after filtering and WGBS > 20X coverage; 60 % CpGs called after filtering) or due to inherent problems with the bisulfite treatment. This corroborates another recent vertebrate study (Lopez-Catalina et al. 2024). Overall, CpG motif methylation sites detected by both methods in the same DNA pool were significantly correlated (Pearson correlation: r=0.80, p-value < 0.001) (Fig 1A). Given that we analysed methylation calls derived from the same DNA pupal pool from *P. californicus* for both ONT sequencing and WGBS, our results demonstrate that ONT technology is a powerful, reliable and maybe even more sensitive tool to determine insect methylomes.

**Fig 1.**
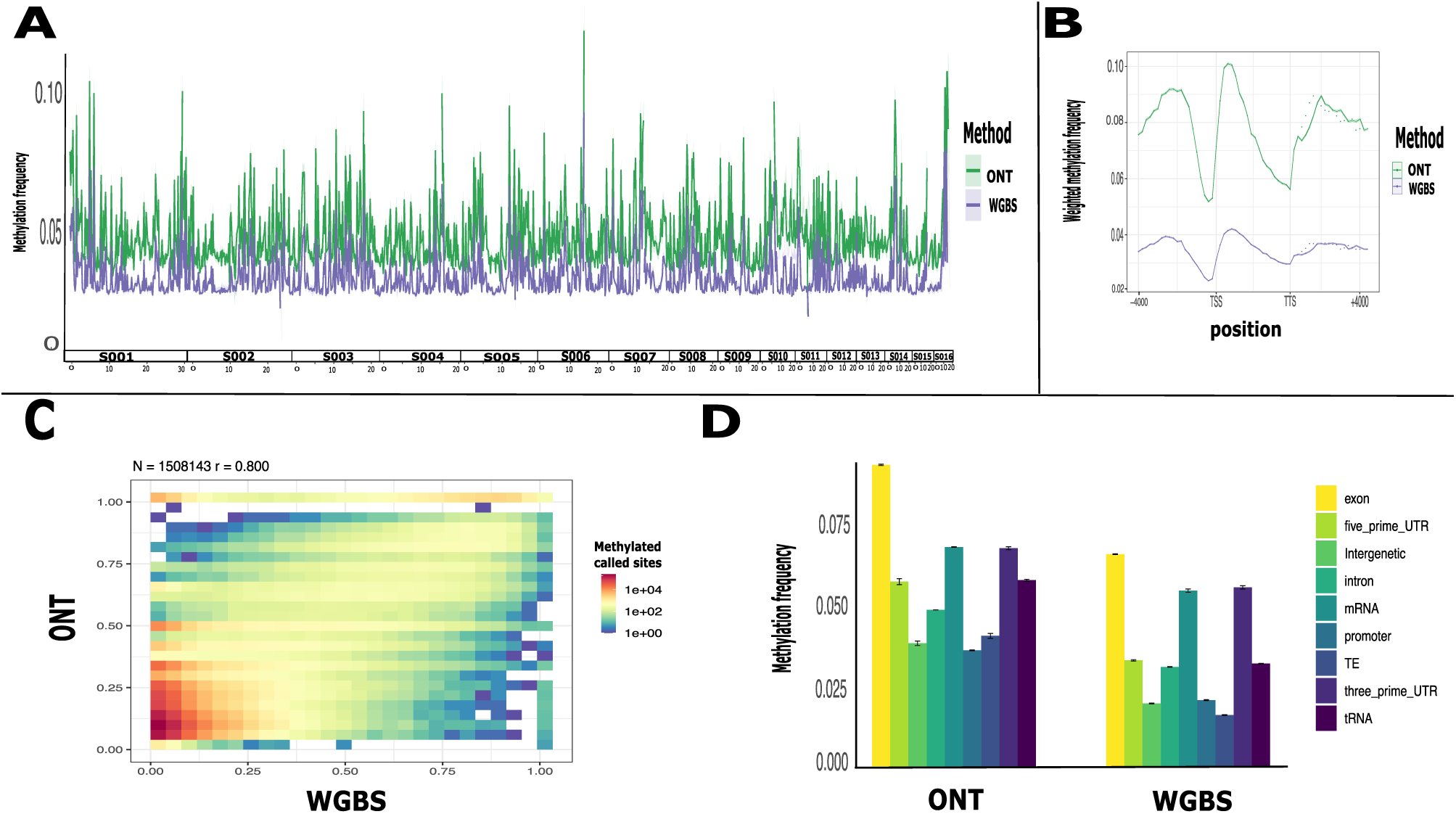
Methylome of *P. californicus* with two sequencing methods Nanopore (ONT) and Whole Genome Bisulfite sequencing (WGBS) using same DNA pupae pools (n=2) comparing the efficiency of both methods to detect CpG methylation rates. A) Average methylation frequency (ONT and WGBS) of two pupae pools along the genome (16 chromosome) B) Weighted frequency methylation average (n=2 pupae pools) (ONT and WGBS) looped over all genes region within a window of 4 kb upstream of the transcription start site (TSS) and a window of 4 kb downstream of the transcription end site (TTS) C) Pearson Correlation of methylation rates of the per-CpG motif detected by both methods (ONT and WGBS), we discard the motif that were only detect by one method to do the correlation analysis D) Average methylation frequency (n=2 pupae pools) (ONT and WGBS) mapped by the genome region annotation.

On average 18.3 million reads per adult sample were generated with ONT sequencing to detect 5mCpG (Table 1). The DNA methylation profile of *P. californicus* was like other Hymenoptera species, characterized by genome wide low levels of CpG methylation (1-10 %). Approximately, 3 % of all CpG sites were methylated (Table 1), with the highest methylation levels observed in genic regions (9.5 %), followed by promoter regions (2.7 %), and intergenic regions, and TEs being least methylated (2.5 %) (Fig 1D). Methylation within genes was higher in exons than in introns, with the second and third exon showing the highest levels of methylation (10%) (Fig 1. B-C) (Fig S3). TEs on the other hand, showed low levels of methylation (2.5 %) (Fig 1D), and we observed a low methylation frequency in TE islands, but this was not significant different when we compared it with the methylation frequency of the rest of the genome (W:395784906, p_value:0.082) (Fig 2. A-B), which could be explained as other regions on the genome as promoters, intergenic regions have low average methylation frequencies. In addition, TE families differed in methylation frequencies. The highest methylation levels were found in “DNA elements” and the lowest in retro and LTR but none was significantly different from the intergenic DNA methylation frequency (Fig S4).

**Fig 2.**
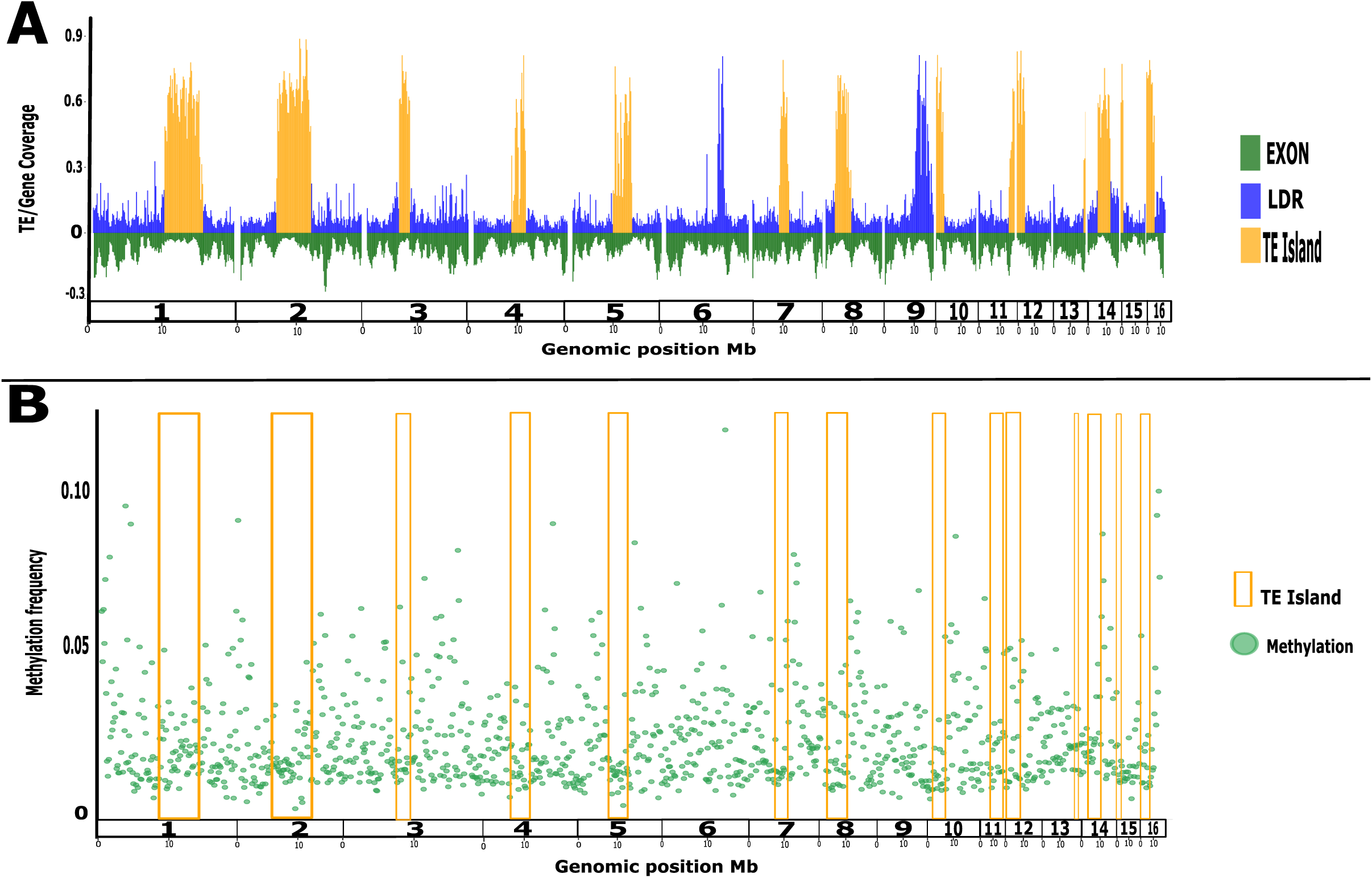
Transposable Elements (TE) annotation and inference of TE islands along *P. californicus* genome. A) Mirror plot of the Genomic positions (Mb) from Long repeat regions (LDR blue color) and TE islands regions (TE yellow color) (above cero line), and the Genome coverage (Exons green color (below cero line) along the genome B) Methylation frequency of CpG methylated sites (green dots) along the genomic regions, yelloe squares represent the TE islands regions.

Genome-wide methylation levels varied among developmental stages and reproductive castes, with larvae showing the lowest percentage of methylated CpG sites (2.9 %), followed by workers (3.1 %), pupae (3.3 %) and highest in queens (3.4 %), however, the overall number of methylated and unmethylated genes did not differ between these samples (Table. 1). Weighted methylation for gene regions was highest for queens and pupae and lowest for worker and larvae (Fig 3. A).

**Fig 3.**
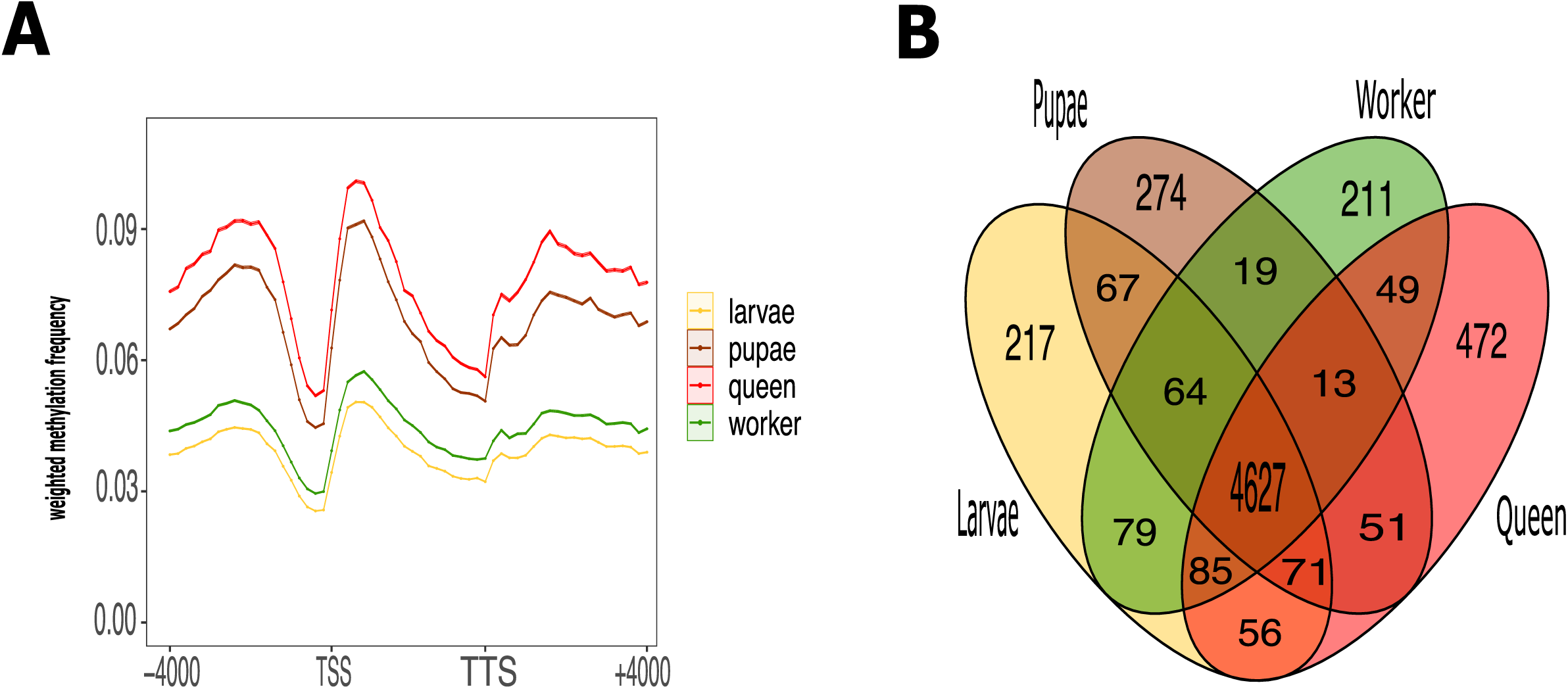
Gene methylation patterns along castes and life stages obtain with the sequencing method ONT A) Weighted frequency methylation average among life stages and castes (n=2 pupae pools, 2 larva pools, 2 worker pools, 10 individuals queens) looped over all genes region within a window of 4 kb upstream of the transcription start site (TSS) and a window of 4 kb downstream of the transcription end site (TTS) B) Venn diagram of Genes body methylated unique and share along stages and castes (larva=yellow, pupae=brown, workers=green, queen=red).

To investigate gene methylation pattern across different castes and life stages, we first computed the average CpG DNA methylation across all genes and performed a binomial test to assess whether coding regions had a significantly higher proportion of methylated cytosines than the genome-wide background level of non-coding regions, which we then called methylated genes (GBM – gene body methylated) in contrast to other genes which were lumped into the un-methylated gene (UM) category (Bewick et al. 2016, Muyle et al. 2021) (see methods above for details). The analyses revealed 5024 methylated genes versus 10,875 unmethylated genes for larvae (Fig 3B, Table. 1, Fig S5), 5186 GBM versus 10,713 UM genes (67%) for workers, 5320 GBM versus 9493 UM, and 5260 GbM versus 10639 UM for queens (Fig 3B, Tab.1, Fig S5). Interestingly, 91% (4698) of the GbM genes were shared between all stages. These shared genes were enriched for functional categories associated with housekeeping processes, including DNA replication, translation and transcription factors, RNA processing and genes corresponding to methyltransferases (Appendix 1).

We found unique GbM genes for each developmental stage (Fig 3B). In larvae, unique GbM genes were associated with lipid metabolism, RNA and DNA synthesis (housekeeping), glycoproteins (immune system) and phosphoynthetases involved in cell growth (Fig 3B, Table S1, Fig S6). In pupae, unique GbM genes were associated with glycosylase activity, metallopeptidase activity, ATP transporters, cell cycle and proteolysis (Fig 3B, Table S2, Fig S7). In workers and queens, unique GbM genes were associated with olfactory receptor activity as well as biosynthetic processes (Fig 3B, Tables S3 and S4, Fig S8-S9).

Promoter regions, defined as a region 200 bp upstream of the TSS, showed the lowest average methylation across all stages. In larvae, we found 1007 Promoter Methylated Genes (PMG) (6.4%) and 14712 Unmethylated Promoter Genes (PUM) (93,6%), while in pupae, there were 1102 PMG (7%) and 14797 PUM (93%), and in adults, we identified 1492 PMG (9%) and 14407 PUM (91%) (Fig S10-S11). We also identified PGM genes unique to each stage. In larva, PMG genes were linked to transcription factors, DNA repair, glucose metabolism, and Krebs cycle (Fig S12). In pupae, PMG genes were associated with DNA replication, transcription and transmembrane transporters (Fig S13). Finally, in adults (workers and queens), PMG genes were enriched in functions related to sensory perception of smell, chemoreception, gustatory receptors and odorant binding (Fig S14).

To study the effect of DNA methylation on gene expression in *P. californicus*, we generated 8-12 million RNA sequencing reads per founding queens (n=22) with an average of 8.5 million reads per sample. After removing low-quality reads, we obtained read counts for 15,948 transcripts. We found that GbM genes showed a higher expression level (FPKM) compared to UM genes (W:29872456 p-value:<2.26e-10 (Fig 4. A), consistent with the role of gene body methylation in the upregulation of gene expression in other insects (Suzuki & Bird 2008, Lyko et al. 2010, Zemach et al. 2010, Bonasio et al. 2012). Additionally, we found a positive correlation between the expression level and methylation frequency of the genes (Pearson-Correlation: r= 0.35 p-value < 2.2e-16, Fig 4. B). Although not significant (W= 6193912, p>0.05), genes with their promoter region methylated showed a slightly higher expression level (Fig S15).

**Fig 4.**
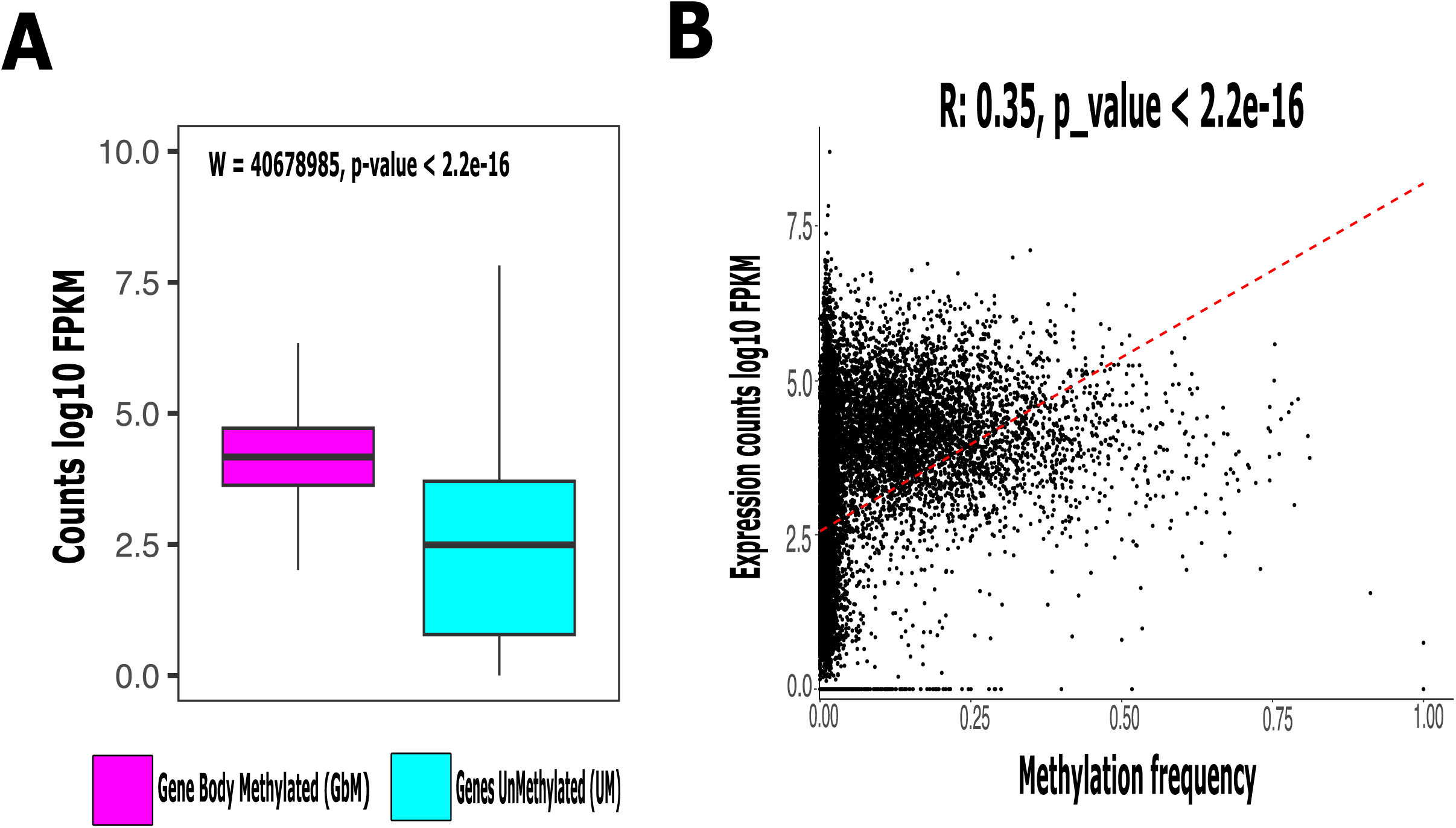
Gene expression level in *P. californicus* queens generated 8-12 million RNA sequencing reads per sample (n=22). A) Expression levels in fragments per kilobase Million (FPKM) were higher in gene body methylated (GbM pink) compared to Unmethylated genes (UM, turquoise) B) Pearson correlation was significantly positive between the expression level (FPKM) and methylation frequency of genes.

Interestingly, but maybe not surprising, gene bodies of the denovo DNA methyltransferase Dnmt3 had a higher methylation frequency for all stages and castes than the maintenance DNA methyltransferases Dnmt1 (Fig S2. A). Indicating its continuous activity in all developmental stages and castes.

## Discussion

This is the first study in ants using ONT sequencing. We found a highly significant correlation between ONT sequencing and the current gold standard in DNA methylation sequencing, WGBS (r= 0,8). This proved that ONT sequencing is well suited for DNA methylation studies in ants.

On average 3% of genome wide CpG sites were methylated, which corroborates results of other Hymenopteran studies (Bewick et al. 2017). Gene regions and in particular gene bodies had the highest methylation frequency. Promoter regions, intergenic regions and transposable elements have low DNA methylation levels (Fig.1, Fig. 2). Furthermore, we could show that gene body methylation significantly increased gene expression levels in *P. californicus* queens (Fig. 4) which is in stark contrast to the function of DNA methylation in vertebrates. Hence, it has been suggested that a reduced methylation frequency of TEs is an epigenetic way “to control and silence TE activity via RNA-directed targeting mechanisms” (Deniz et al. 2019, Perez et al. 2023), just in the opposite way as in vertebrates where an increase in DNA methylation reduces TE expression, but the concrete mechanism is unknown. Nonetheless, methylation frequency of TEs varies across species in invertebrates, hence it needs further detailed studies to clarify the effect of DNA methylation on TEs expression in invertebrates (Lyko et al. 2010, de Mendoza et al. 2020, Lewis et al. 2020, Perez et al. 2023).

Developmental stages show distinct methylation frequencies which confirms its role as a proximate mechanism for development and may affect caste determination in social insects (Yan et al. 2015, Kozeretska, et al. 2017, Morandin et al. 2019). In our study DNA methylation levels in larvae were lower which could be explained in part by the fact that most of the DNA methylation marks are removed during gametogenesis and reset during development (Smallwood & Kelsey 2012). However, it might have other functions related to juvenile stages because several studies found lower DNA methylation levels in juvenile life stages of both vertebrates and invertebrates (Wang & Bhandari 2019, Planques et al. 2021, Perez et al. 2023). Furthermore, juvenile stages (larvae and pupae), in social insects, show in general less morphological, physiological and behavioral variation than adults and might therefore need fewer genes to be expressed (Morandin et al. 2014, Harrison et al. 2015). Schrader et al. 2014 found for example that queens expressed significantly more genes than larvae in TE islands of the ant *Cardiocondyla obscurior*.

We discovered many differences in DNA methylation especially at the level of gene body methylation between developmental stages (larvae, pupae, adults) and castes (workers and queens) in *P. californicus*. Our results coincide with other studies in ants, which also found gene expression differences between developmental stages and castes, expressing specific genes related with important functions for their respective life stage or caste (Gstöttl et al. 2020). In *P. californicus* GbM genes in larvae were enriched for lipid metabolism and storage, glycoproteins (immune system) and phosphosynthetases (cell grow); while for pupae GbM genes were enriched for glycosylase activity and cell cycle (cell grow). Morandin et al. (2014) and Gstöttl et al. (2020) also found that larvae and pupae upregulated genes involved in growth and tissue buildup. Similarly for several other social insect species, Kapheim et al. (2017) documented that larvae have upregulated genes related to lipid storage and lipid metabolism.

For queens and workers, we found that many differentially methylated genes between workers and queens were enriched for categories involved with chemical communication, e.g. chemoreception, gustatory receptors and odorant binding. These categories were also enriched in a transcriptomic study on *P. californicus* founding queens (Helmkampf et al. 2016). Chemical communication is for social insects very essential across many different functions like nestmate discrimination or caste recognition. Furthermore, queens also have uniquely methylated genes involved in metabolic process, DNA damage checkpoint, and metalloproteases which are involved in modulation of cell growth, inflammation, immunity, and hormone processing. This coincides with transcriptomic studies which have found that queens upregulated DNA repair genes (Gstöttl et al. 2020) which has been associated with anti-aging strategies (Lucas, Privman & Keller, 2016, Gstöttl et al. 2020). Finally, we found one methylated gene related to the neuropeptide corazonin, which has been reported as a key regulator of caste identity in ants and other social insects (Gosposic et al. 2017). Nonetheless, we should be aware that some of the variation of the methylated genes between workers and queens could be due to population differences, as worker samples were derived from lab colonies of queens collected in California and queen samples were collected from populations in Arizona.

Gene body methylated genes shared between all stages are enriched for functional categories associated with housekeeping functions such as DNA and RNA processing, cell cycle, mitochondrial respiration, HsP30 genes and several methyltransferases (Appendix 1). This corroborates other studies which found that mean methylation frequency (methylation level) of genes is highest in functional categories associated with housekeeping functions (Perez et al. 2019). Furthermore, up-regulation of genes linked to “Regulation of Gene Expression and Epigenetic Mechanisms”, has been related to a high fecundity and a long life in social insects (Gstöttl et al. 2020).

The methylation of the promoter region has also been associated with an increase of gene expression in insects (Bewick et al. 2017). In the harvester ant *P. californicus* we found a trend but no significant relationship between promoter methylation and higher expression. Some studies have suggested that promoter methylation plays a supporting role for other gene expression mechanisms, as they also found no evidence of promoter methylation and gene expression or silencing (Keller et al. 2016, Perez et al. 2023).

Dnmt1 and Dnmt3 were both methylated in *P. californicus,* although Dnmt3 showed a much higher DNA methylation frequency (Fig S2). Interestingly in *P. californicus*, both Dnmts have diverged between populations and social forms (Errbii et al. 2024). Since genetic variation can affect DNA methylation of a gene and its expression, these genetic differences might be linked via differences in DNA methylation to the observed social polymorphism (Wedd et al. 2016). Dnmt1 1 is crucial for maintaining the DNA methylation patterns inherited during insect development and is also involved in regulating gene expression that contribute to division of labour between reproductive (queens) and non-reproductive individuals (workers) in honeybees (Kurcharski et al. 2009, Lyko et al. 2010, Liebrecht et al. 2016). However, Dnmt1 gene knockout experiments have shown that normal somatic gene expression in plants and insects does not need gene body methylation (Bewick et al. 2019). Similar experimental approaches are needed in ants, to determine if DNA methylation has a direct epigenetic control on gene expression and caste determination in ants.

## Conclusion

The harvester ant *P. californicus* like other solitary and social Hymenoptera (bumble bees, honeybees and the parasitoid wasp *Nasonia*) have a small but significant proportion of their CpGs methylated. DNA methylation is like in other invertebrates highly concentrated in exons, and gene expression in *P. californicus* increases with gene body methylation. We could show that DNA methylation varies between developmental stages, lowest in larvae, followed by adult workers and pupae, and castes, with queens having more methylated sites than workers. We found a significant number of caste and stage specific methylated genes. Especially, interesting are differences in DNA methylation between workers and queens in genes related to chemical perception and recognition. These findings warrant further in-depth studies of the link between DNA methylation, gene expression and phenotypic plasticity in *P. californicus* to understand the role of epigenetic modifications for phenotypic variation, development, caste determination and social polymorphism in general and *P. californicus* in particular.

## Supporting information

Supplementary Information including Figures and Tables

## Acknowledgments

Hilde Schwitte performed all the DNA extractions. Kathrin Brüggemann generated the MinION libraries which produced ONT sequences. Bernice Speers and Aline Muyle for generated the R scripts for some methylation analysis used in this manuscript.

## Author contributions

M.E. contributed by generating the genome annotation (Erbii et al. unpublished), data analysis, collected queens for transcriptome data and manuscript editing. J.R contribute with data analysis and manuscript editing. L.S contributed with data analysis and manuscript editing. J.G. conceived the project proposal, collected queens for transcriptome data and for develop the colonies at the lab, contributed to manuscript writing and editing. T.Ch.P. collected the founding queens for the methylome analysis, analysed most of the data and wrote the manuscript.

## Data availability statement

The transcriptome data and methylation calls have been deposited with links to BioProject accession no. **BioProject:PRJNA1234538** in the NCBI BioProject database (https://dataview.ncbi.nlm.nih.gov/object/PRJNA1234538), and they will be released when the manuscript is published in a peer-review journal. The required scripts to do the methylation analysis (e.g. basecalling, methylation calling, bedtools, samtools, and R script) are available in this repositor https://github.com/TaniaChP79/P.cal-methylome

## Notes

### Competing Interest Statement

The authors have declared no competing interest.

