## Supplementary Information including Figures and Tables for "Castes and developmental stages of the harvester ant *P. californicus* differ in genome-wide and gene specific DNA methylation"

This file includes:

Figs. S1- S15

Tables S1-S4

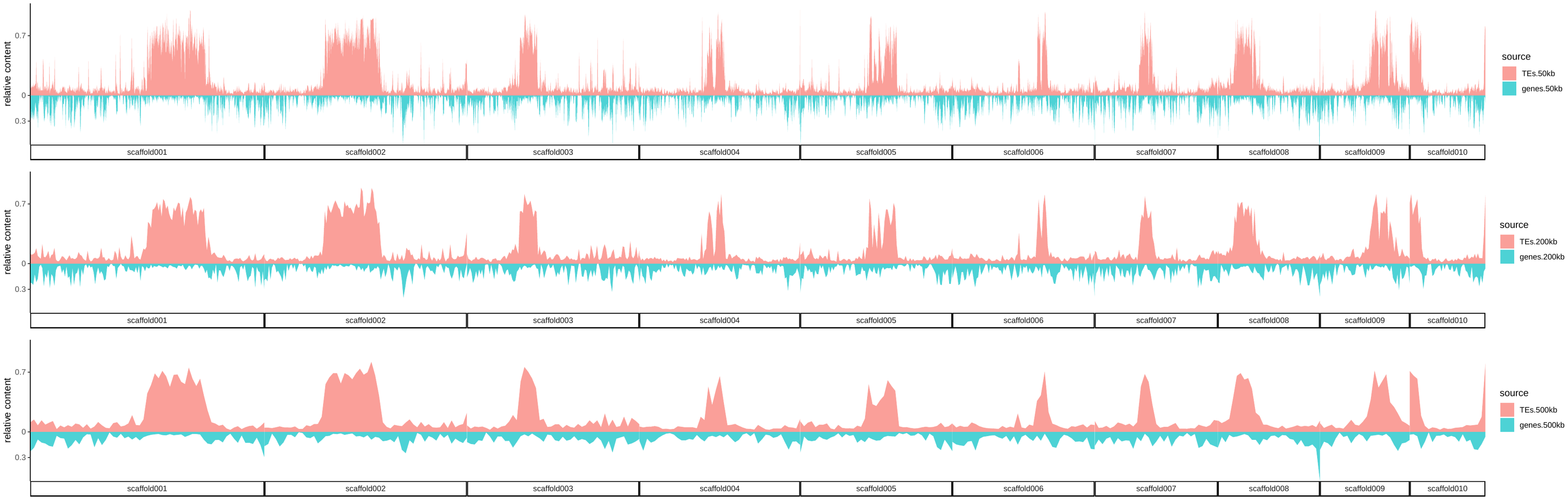

S1. Relative content of Transposable Elements (TEs pink) and Genes (turquoise) along 10 scaffolds (chromosomes) using a 50 kb long genomic window along the genome of *P. californicus*.

S2. DNMT1 and DNMT3 methylation frequency in *P. californicus* and phylogenetic similarity with close related species. A) Average methylation frequency (%) of DNMT1 and DNMT3 along life stages (n=2 pupae pools, 2 larva pools, adults= 2 worker pools + 10 individuals queens) B) DNMT1 maximum likelihood phylogenetic tree along some related Hymenopteran species with DNMT1 sequences available 3) DNMT3b maximum likelihood phylogenetic tree along some related Hymenopteran species with DNMT3b sequences available

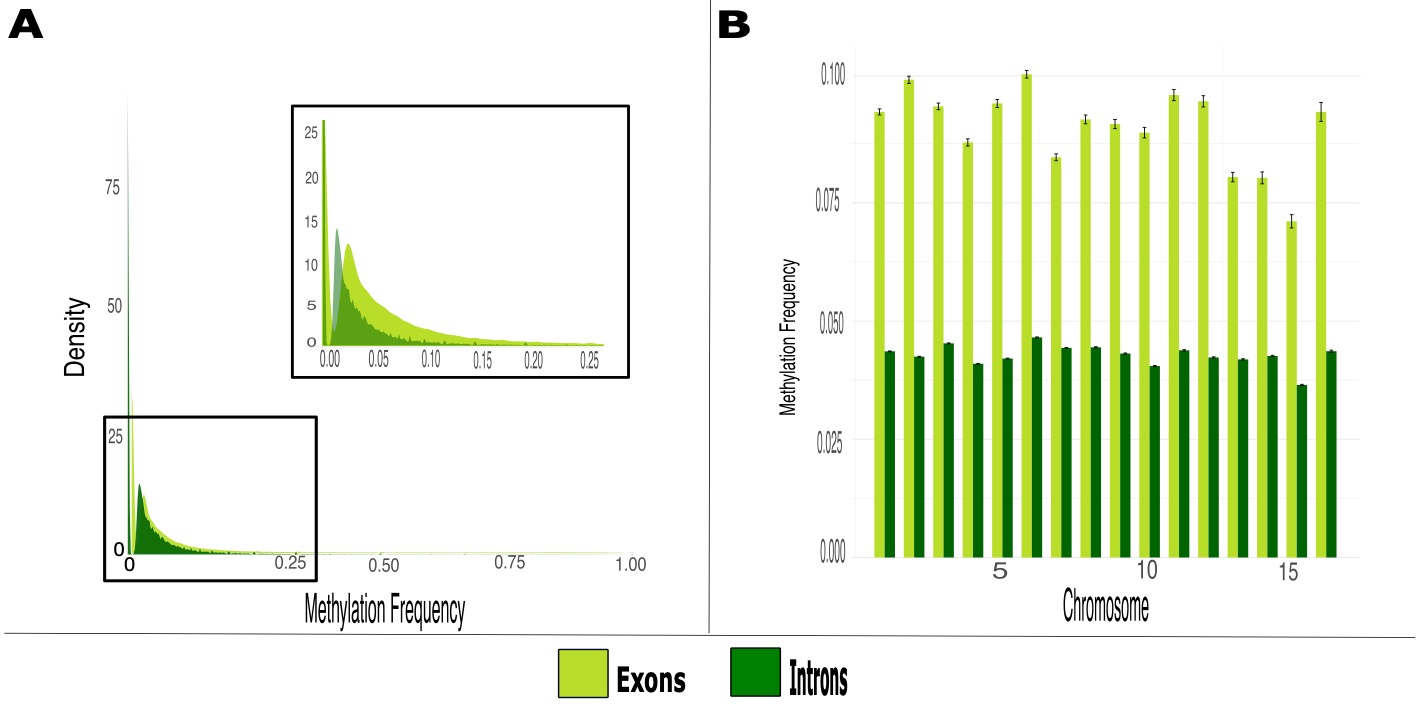

S3. Average Methylation frequency of exons and introns along *P. californicus* genome. A) Methylation frequency of exons and intros according with the CpG Density, on the x-axis we plot the methylation frequency calculated with ONT and on the y-axis we plotted the density (CpG methylated sites / total CpG sites) on percentages (Exons=dark green, Introns=light green) B) Methylation frequency of exons and introns along the 16 chromosomes.

S4. Average Methylation frequency and standard deviation of Transposable Elements (TEs) families in *P. californicus* genomes.

S5. Gene methylation patterns along life stages obtain with the sequencing method ONT A) Venn diagram of Genes body methylated unique and share along stages and castes (larva=yellow, pupae=brown, adults=red) (n=2 pupae pools, 2 larva pools, adults= 2 worker pools + 10 individuals queens) B) Venn diagram of Genes body Unmethylated unique and share along stages and castes (larva=yellow, pupae=brown, adults=red) (n=2 pupae pools, 2 larva pools, adults= 2 worker pools + 10 individuals queens).

S6. GO enrichment analysis for all genes body methylated unique to larvae(n=2 larvae pools) with significant gene-body methylation with a p-value < 0.05 representing significantly enriched GO terms.

S7. GO enrichment analysis for all genes body methylated unique to pupae (n=2 pupae pooles) with significant gene-body methylation with a p-value < 0.05 representing significantly enriched GO terms.

S8. GO enrichment analysis for all genes body methylated unique to queen (n=10 individual queens) with significant gene-body methylation with a p-value < 0.05 representing significantly enriched GO terms.S9. GO enrichment analysis for all genes body methylated unique to workers (n=2 worker pools) with significant gene-body methylation with a p-value < 0.05 representing significantly enriched GO terms.

S10. Promoter methylated genes patterns along life stages obtain with the sequencing method ONT A) Venn diagram of Promoter methylated genes unique and share along stages and castes (larva=yellow, pupae=brown, adults=red) (n=2 pupae pools, 2 larva pools, adults= 2 worker pools + 10 individuals queens) B) Venn diagram of Promoter Unmethylated genes unique and share along stages and castes (larva=yellow, pupae=brown, adults=red) (n=2 pupae pools, 2 larva pools, adults= 2 worker pools + 10 individuals queens).

S11. GO enrichment analysis for all genes with significant promoter methylation common to all life stages n=2 pupae pools, 2 larva pools, adults= 2 worker pools + 10 individuals queens) p-value < 0.05 representing significantly enriched GO terms.

S12. GO enrichment analysis for all genes unique to larvae with significant promoter methylation (n=2 pupae pools) p-value < 0.05 representing significantly enriched GO terms.

S12. GO enrichment analysis for all genes unique to larvae with significant promoter methylation (n=2 pupae pools) p-value < 0.05 representing significantly enriched GO terms.

S13. GO enrichment analysis for all genes unique to pupae with significant promoter methylation (n=2 pupae pools) p-value < 0.05 representing significantly enriched GO terms.

S14. GO enrichment analysis for all genes unique to adults with significant promoter methylation (adults= 2 worker pools + 10 individuals queens) p-value < 0.05 representing significantly enriched GO terms.

S15. Gene expression level of genes with promoter methylation in *P. californicus* queens generated 8-12 million RNA sequencing reads per sample (n=22). Expression levels in fragments per kilobase Million (FPKM) were non-significant higher in gene with promoter methylation (pink) compared to Unmethylated promoter genes (turquoise).

Table S1. Gene differential methylated unique for larvae significant associated to Ontology using a Fisher test. GO.ID (GO Term Identification number), NumDMGs (number differential body methylated genes), pvalue (Fisher test).

| GO.ID | GOTerm | NumDMGs | Pvalue | Ontology |
| --- | --- | --- | --- | --- |
| GO:0008442 | 3-hydroxyisobutyrate dehydrogenase activ... | 1 | 0.012 | Molecular_Function |
| GO:0004329 | formate-tetrahydrofolate ligase activity | 1 | 0.012 | Molecular_Function |
| GO:0004677 | DNA-dependent protein kinase activity | 1 | 0.012 | Molecular_Function |
| GO:0016316 | phosphatidylinositol-3.4-bisphosphate 4-... | 1 | 0.012 | Molecular_Function |
| GO:0008681 | 2-octaprenyl-6-methoxyphenol hydroxylase... | 1 | 0.012 | Molecular_Function |
| GO:0033925 | mannosyl-glycoprotein endo-beta-N-acetyl... | 1 | 0.012 | Molecular_Function |
| GO:0004029 | aldehyde dehydrogenase (NAD+) activity | 1 | 0.012 | Molecular_Function |
| GO:0004334 | fumarylacetoacetase activity | 1 | 0.012 | Molecular_Function |
| GO:0004642 | phosphoribosylformylglycinamidine syntha... | 1 | 0.012 | Molecular_Function |
| GO:0017017 | MAP kinase tyrosine/serine/threonine pho... | 1 | 0.012 | Molecular_Function |
| GO:0004531 | deoxyribonuclease II activity | 1 | 0.012 | Molecular_Function |
| GO:0004372 | glycine hydroxymethyltransferase activit... | 1 | 0.012 | Molecular_Function |
| GO:0005185 | neurohypophyseal hormone activity | 1 | 0.012 | Molecular_Function |
| GO:0003796 | lysozyme activity | 2 | 0.023 | Molecular_Function |
| GO:0004459 | L-lactate dehydrogenase activity | 2 | 0.023 | Molecular_Function |
| GO:0004488 | methylenetetrahydrofolate dehydrogenase ... | 2 | 0.023 | Molecular_Function |
| GO:0004420 | hydroxymethylglutaryl-CoA reductase (NAD... | 2 | 0.023 | Molecular_Function |
| GO:0045296 | cadherin binding | 4 | 0.046 | Molecular_Function |

Table S1. Continued.

| GO.ID | GOTerm | NumDMGs | Pvalue | Ontology |
| --- | --- | --- | --- | --- |
| GO:0035735 | intraciliary transport involved in ciliu... | 1 | 0.0084 | Biological_Process |
| GO:0019264 | glycine biosynthetic process from serine | 1 | 0.0084 | Biological_Process |
| GO:0035999 | tetrahydrofolate interconversion | 1 | 0.0084 | Biological_Process |
| GO:0010960 | magnesium ion homeostasis | 1 | 0.0084 | Biological_Process |
| GO:0061780 | mitotic cohesin loading | 1 | 0.0084 | Biological_Process |
| GO:0032958 | inositol phosphate biosynthetic process | 2 | 0.0168 | Biological_Process |
| GO:0035721 | intraciliary retrograde transport | 2 | 0.0168 | Biological_Process |
| GO:0006189 | 'de novo' IMP biosynthetic process | 3 | 0.0251 | Biological_Process |
| GO:0006303 | double-strand break repair via nonhomolo... | 3 | 0.0251 | Biological_Process |
| GO:0031145 | anaphase-promoting complex-dependent cat... | 4 | 0.0333 | Biological_Process |
| GO:0031463 | Cul3-RING ubiquitin ligase complex | 1 | 0.0086 | Cellular_Component |
| GO:0005868 | cytoplasmic dynein complex | 5 | 0.0423 | Cellular_Component |

Table S2. Gene differential methylated unique for pupae significant associated to Ontology using a Fisher test. GO.ID (GO Term Identification number), NumDMGs (number methylated genes), pvalue (Fisher test).

| GO.ID | GOTerm | NumDMGs | pvalue | Ontology |
| --- | --- | --- | --- | --- |
| GO:0008237 | metallopeptidase activity | 293 | 0.00099 | Molecular Function |
| GO:0008173 | RNA methyltransferase activity | 36 | 0.00177 | Molecular Function |
| GO:0008270 | zinc ion binding | 537 | 0.00184 | Molecular Function |
| GO:0004181 | metallocarboxypeptidase activity | 19 | 0.00248 | Molecular Function |

Table S2. Continued.

| GO.ID | GOTerm | NumDMGs | Pvalue | Ontology |
| --- | --- | --- | --- | --- |
| GO:0004714 | transmembrane receptor protein tyrosine ... | 8 | 0.0056 | Molecular Function |
| GO:0015293 | symporter activity | 12 | 0.01271 | Molecular Function |
| GO:0004222 | metalloendopeptidase activity | 140 | 0.01639 | Molecular Function |
| GO:0019843 | rRNA binding | 16 | 0.02225 | Molecular Function |
| GO:0005471 | ATP:ADP antiporter activity | 2 | 0.02904 | Molecular Function |
| GO:0042132 | fructose 1.6-bisphosphate 1-phosphatase ... | 3 | 0.04325 | Molecular Function |
| GO:0003905 | alkylbase DNA N-glycosylase activity | 3 | 0.04325 | Molecular Function |
| GO:0006508 | Proteolysis | 499 | 0.014 | Biological Process |
| GO:0051013 | microtubule severing | 2 | 0.029 | Biological Process |
| GO:0034975 | protein folding in endoplasmic reticulum | 2 | 0.029 | Biological Process |
| GO:1990544 | mitochondrial ATP transmembrane transpor... | 2 | 0.029 | Biological Process |
| GO:0007052 | mitotic spindle organization | 2 | 0.029 | Biological Process |
| GO:0140021 | mitochondrial ADP transmembrane transpor... | 2 | 0.029 | Biological Process |
| GO:0035060 | brahma complex | 1 | 0.012 | Celular component |
| GO:0008352 | katanin complex | 1 | 0.012 | Celular component |
| GO:0005743 | mitochondrial inner membrane | 51 | 0.019 | Celular component |
| GO:0031209 | SCAR complex | 2 | 0.024 | Celular component |
| GO:0071339 | MLL1 complex | 4 | 0.048 | Celular component |
| GO:0089701 | U2AF complex | 4 | 0.048 | Celular component |

Table S3. Gene differential methylated unique for worker significant associated to Ontology using a Fisher test. GO.ID (GO Term Identification number), NumDMGs (number methylated genes), pvalue (Fisher test).

| GO.ID | GOTerm | NumDMGs | pvalue | Ontology |
| --- | --- | --- | --- | --- |
| GO:0005506 | iron ion binding | 116 | 0.00021 | Molecular_Function |
| GO:0016705 | oxidoreductase activity. acting on paire... | 127 | 0.00036 | Molecular_Function |
| GO:0004497 | monooxygenase activity | 110 | 0.00102 | Molecular_Function |
| GO:0020037 | heme binding | 121 | 0.00167 | Molecular_Function |
| GO:0008449 | N-acetylglucosamine-6-sulfatase activity | 1 | 0.01057 | Molecular_Function |
| GO:0016263 | glycoprotein-N-acetylgalactosamine 3-bet... | 1 | 0.01057 | Molecular_Function |
| GO:0000285 | 1-phosphatidylinositol-3-phosphate 5-kin... | 1 | 0.01057 | Molecular_Function |
| GO:0004047 | aminomethyltransferase activity | 1 | 0.01057 | Molecular_Function |
| GO:0015016 | [heparan sulfate]-glucosamine N-sulfotra... | 2 | 0.02103 | Molecular_Function |
| GO:0016803 | ether hydrolase activity | 3 | 0.03139 | Molecular_Function |
| GO:0004861 | cyclin-dependent protein serine/threonin... | 3 | 0.03139 | Molecular_Function |
| GO:0031418 | L-ascorbic acid binding | 4 | 0.04163 | Molecular_Function |
| GO:0004984 | olfactory receptor activity | 313 | 0.04651 | Molecular_Function |
| GO:0005096 | GTPase activator activity | 34 | 0.04976 | Molecular_Function |
| GO:0009890 | negative regulation of biosynthetic proc... | 64 | 0.01 | Biological_Process |
| GO:0032963 | collagen metabolic process | 1 | 0.01 | Biological_Process |
| GO:0048208 | COPII vesicle coating | 1 | 0.01 | Biological_Process |
| GO:0016266 | O-glycan processing | 1 | 0.01 | Biological_Process |
| GO:0006103 | 2-oxoglutarate metabolic process | 1 | 0.01 | Biological_Process |
| GO:0006914 | Autophagy | 23 | 0.022 | Biological_Process |
| GO:0005096 | GTPase activator activity | 34 | 0.04976 | Molecular_Function |

Table S3. Continued.

| GO.ID | GOTerm | NumDMGs | pvalue | Ontology |
| --- | --- | --- | --- | --- |
| GO:0009058 | biosynthetic process | 1442 | 0.025 | Biological_Process |
| GO:0006546 | glycine catabolic process | 3 | 0.03 | Biological_Process |
| GO:0007608 | sensory perception of smell | 313 | 0.036 | Biological_Process |
| GO:0019722 | calcium-mediated signaling | 4 | 0.04 | Biological_Process |
| GO:0006493 | protein O-linked glycosylation | 6 | 0.048 | Biological_Process |
| GO:0009353 | mitochondrial oxoglutarate dehydrogenase... | 1 | 0.0078 | Cellular_Component |
| GO:0055087 | Ski complex | 3 | 0.0232 | Cellular_Component |
| GO:0008278 | cohesin complex | 3 | 0.0232 | Cellular_Component |
| GO:0016020 | Membrane | 968 | 0.0357 | Cellular_Component |
| GO:0045211 | postsynaptic membrane | 6 | 0.0458 | Cellular_Component |

Table S4. Gene differential methylated unique for queen significant associated to Ontology using a Fisher test. GO.ID (GO Term Identification number), NumDMGs (number methylated genes), pvalue (Fisher test).

| GO.ID | GOTerm | NumDMGs | Pvalue | Ontology |
| --- | --- | --- | --- | --- |
| GO:0005549 | odorant binding | 331 | 5e-11 | Molecular_Function |
| GO:0004984 | olfactory receptor activity | 313 | 1.4e-09 | Molecular_Function |
| GO:0008237 | metallopeptidase activity | 293 | 7.9e-08 | Molecular_Function |
| GO:0008270 | zinc ion binding | 537 | 4.3e-06 | Molecular_Function |
| GO:0031177 | phosphopantetheine binding | 10 | 0.0015 | Molecular_Function |
| GO:0015018 | galactosylgalactosylxylosylprotein 3-bet... | 3 | 0.0017 | Molecular_Function |
| GO:0008061 | chitin binding | 57 | 0.0025 | Molecular_Function |

Table S4. Continued.

| GO.ID | GOTerm | NumDMGs | Pvalue | Ontology |
| --- | --- | --- | --- | --- |
| GO:0005506 | iron ion binding | 116 | 0.0226 | Molecular_Function |
| GO:0030955 | potassium ion binding | 1 | 0.0243 | Molecular_Function |
| GO:0050113 | inositol oxygenase activity | 1 | 0.0243 | Molecular_Function |
| GO:0004743 | pyruvate kinase activity | 1 | 0.0243 | Molecular_Function |
| GO:0008200 | ion channel inhibitor activity | 1 | 0.0243 | Molecular_Function |
| GO:0071858 | corazonin receptor binding | 1 | 0.0243 | Molecular_Function |
| GO:0004315 | 3-oxoacyl-[acyl-carrier-protein] synthas... | 12 | 0.0331 | Molecular_Function |
| GO:0004496 | mevalonate kinase activity | 30 | 0.0355 | Molecular_Function |
| GO:0004497 | monooxygenase activity | 110 | 0.0374 | Molecular_Function |
| GO:0004605 | phosphatidate cytidylyltransferase activ... | 2 | 0.0481 | Molecular_Function |
| GO:0007608 | sensory perception of smell | 313 | 1.5e-10 | Biological_Process |
| GO:0050909 | sensory perception of taste | 55 | 3.2e-06 | Biological_Process |
| GO:0006508 | Proteolysis | 499 | 1.3e-05 | Biological_Process |
| GO:0045823 | positive regulation of heart contraction | 1 | 0.023 | Biological_Process |
| GO:0019310 | inositol catabolic process | 1 | 0.023 | Biological_Process |
| GO:0016020 | Membrane | 968 | 0.00014 | Cellular_Component |
| GO:0031012 | extracellular matrix | 82 | 0.01295 | Cellular_Component |
| GO:0005576 | extracellular region | 105 | 0.0171 | Cellular_Component |
| GO:0036064 | ciliary basal body | 2 | 0.04775 | Cellular_Component |
